# Biofilm Associated Genotypes of Multidrug-Resistant *Pseudomonas aeruginosa*

**DOI:** 10.1101/713453

**Authors:** J. Redfern, J. Wallace, A. van Belkum, M. Jaillard, E. Whittard, R. Ragupathy, J. Verran, P. Kelly, M.C. Enright

## Abstract

*Pseudomonas aeruginosa* is a ubiquitous environmental microorganism that is also a common cause of nosocomial infections that vary in severity from chronic wound infections to pneumonia, bloodstream infections and sepsis. Its ability to survive in many different environments and persistently colonize human tissue is linked to its presence within biofilms that form on indwelling device surfaces such as plastics and stainless steel. Biofilm promotes bacterial adhesion and survival on surfaces, reduces susceptibility to desiccation, and the actions of antibiotics and disinfectants. Recent genome sequencing studies demonstrate that *P. aeruginosa* is a highly diverse species with a very large pan-genome consistent with its adaptability to differing environments. However, most MDR infections are caused by a small number of “high-risk” clones or lineages that recently emerged and spread globally.

In our 2017 study of the resistome of *P. aeruginosa* we confirmed the power of genome-wide association (GWAS) techniques to explore the genetic basis of several antibiotic resistance phenotypes and discovered 46 novel putative resistance polymorphisms. In this study we sought to examine genetic associations within a subset of these isolates with simple biofilm phenotypes. We examined the genetic basis for biofilm production on polystyrene at room temperature (22°C) and body temperature (37°C) within a total of 280 isolates. 69% of isolates (*n*=193) produced more biofilm mass at 22°C, whilst those producing more biofilm at 37°C had reduced optical density _540_ variation. We found statistically significant associations with *lpxO* and other genes associated with arsenic resistance to be significantly associated with this trait. *lpxO* which encodes a lipid A hydroxylase and arsenic reduction genes have previously been found to be associated with biofilm production in this species. We analyzed 260 ST111 and ST235 genomes and found considerable genetic variation between isolates in their content of genes previously found associated with biofilm production. This is indicative of a highly variable and flexible population within these clades with frequent emergence of successful sub-lineages. Analysis of 48 of these isolates’ ability to form biofilm on stainless steel surfaces showed that a ‘good’ biofilm-forming phenotype had significant intra-clone variation, independent of core genome phylogeny with pan-genome analysis, suggesting a possible association and involvement of components of the type IV secretion system. However, GWAS and pan-GWAS analyses yielded weaker statistical significance. This study confirms GWAS and pan-GWAS trait associations can be performed for biofilm phenotype and produce data in agreement with each other. This panel of 280 study isolates, matched to genomic data has potential for the investigation of other phenotypes in *P. aeruginosa* perhaps as part of a growing database / collection. A representative, curated, genome sequenced collection should increase in usefulness as it grows offering increasing statistical power.

**Importance:** *P. aeruginosa* is a major cause of multiply antibiotic infections worldwide but it is also found in many hospital and natural environments, especially aquatic ones. In this study we examined genetic polymorphism associated with biofilm production at room temperature and at body temperature, the biofilm associated gene repertoire of two major MDR clones and also genetic polymorphisms associated with biofilm production on stainless steel. Using these genome-wide and pan-genome wide association methods we identified / confirmed potential key genes involved in biofilm production and survival of *P. aeruginosa*. The study demonstrates the potential usefulness of large, genome sequenced isolate collections such as ours, to better understand the genetics underlying phenotypic diversity in this species.

## Introduction

*Pseudomonas aeruginosa* is a mono-flagellate, Gram-negative bacterium that is present in most environments. A frequent coloniser of humans, other animals and plants, it also is a very common opportunistic pathogen able to grow at a variety of temperatures^1^. It is a leading cause of severe, life-threatening nosocomial human infections and the major pathogen associated with lung infections of patients with cystic fibrosis. It is has been classified as one of the major species associated with multiple antimicrobial resistance of urgent public health concern by the Infectious Diseases Society of America^2^, Centers for Disease Control and Prevention and the World Health Organization.

A variety of typing methods including multilocus sequence typing (MLST)^3^ and more recently, genomic sequencing studies^4–7^ have shown that *P. aeruginosa* has a non-clonal, epidemic population structure in which successful clones occasionally arise and are globally transmitted^8^. Multidrug resistant (MDR) isolates of clones associated with outbreaks of infection belonging to MLST Sequence Type (ST) 111 and ST235 are two such ‘high risk’ clones that have global distributions and numerous and transferable antibiotic resistances^9^. Isolates of these clones contain a large number of horizontally transferred β-lactamase genes^4^ and are highly virulent in comparison to a third, less significant clone – ST175^10^.

The antibiotic resistome of both ST111 and ST235 clones have been characterised in attempts to understand relationships between antimicrobial resistance (AMR) phenotype and genotype. A genomic analysis of 87 UK isolates belonging to ST111 demonstrated divergence from a common ancestor estimated to have emerged in approximately 1965, with the majority of isolates carrying the β-lactam gene VIM-2^5^. A similar analysis of 79 ST235 isolates predicted a more recently evolved clone, the ancestral genotype of which emerged around 1984, coinciding with the widespread use of fluoroquinolones for the treatment of gram-negative infections^11^.

Whilst understanding the relationship between a pathogen’s genome, epidemiological surveillance, clinical treatment and prevention success is critically important^12,13^, *P. aeruginosa* has many phenotypic features that also promote its success as a pathogen. In common with other bacterial pathogens *P. aeruginosa* usually exists within biofilms. These are complex microbial communities associated with extracellular polymeric matrices that help bacteria resist desiccation, mechanical removal and the actions of antibiotics and disinfectants^14^. In human tissue, biofilm-associated *P. aeruginosa* are difficult to eradicate and represent infectious foci that can lead to serious systemic disease. *P. aeruginosa* biofilms are found at surface-liquid, surface-air and liquid-air interfaces, and are a significant clinical problem in wounds and the cystic fibrosis lung^15^. Polymicrobial biofilms are also recognized as important niches for MDR evolution, as they represent a significant reservoir for horizontal gene transfer within and between bacterial species^16^.

In addition to comprehending *in vivo* biofilm formation, understanding *P. aeruginosa* biofilm development and persistence on other surfaces is required. Stainless steel is the most common surface material used in many industries where control of microorganisms is important, including healthcare. Properties of stainless steel include resistance to corrosion, easiness to clean and a level of hardness likely to limit scratches and other defects^17^. However, many of these surfaces have been shown to allow the formation of biofilms, for example in hard-to-clean locations, and those with a favourable environment, such as sinks and pipes ^18^.

Both ST111 and ST235 isolates have been shown to produce significantly more biofilm compared to those from non-high-risk clones^19^ whilst being over-represented in locations such as hospital sink pipes^20^. A recent study strongly implicates the presence of *P. aeruginosa* biofilm in hospital waste-water pipes as being responsible for an outbreak of infection due to MDR ST111 and ST235 in a haematology unit^21^. A policy promoting replacement of sink units to reduce such sources of infection has now been rolled out in the United Kingdom.

There is a need to better understand the clinical importance of biofilms in hospital environments starting with how biofilm production changes when *P. aeruginosa* cells move from the hospital environment to the human body. Genome Wide Association Studies (GWAS) have been used to great effect in identifying genetic determinants contributing to disease pathology in human medicine. Thus far, microbial GWAS and more generic Whole Genome Sequencing (WGS) studies have focused on the molecular epidemiology of infectious disease and particularly on associations between the genome and antimicrobial resistance and / or pathogenicity. There is huge potential for such methods to help understand the genetic basis of other phenotypes, such as key processes that are important in microbial ecology or industrial microbiology^22,23^.

In this study we examine the associations between aspects of the *P. aeruginosa* pan-genomes and their biofilm-producing activity at different temperatures or on stainless steel and examine the genetic attributes associated with biofilm production in MDR clones ST111 and ST235

## Methods

### Bacterial isolates and genome sequences

This study analyzed two different sets of *P. aeruginosa* isolates. The first set, here-on-in described as ‘temperature-dependent biofilm analysis’ contained 280 isolates (Supp Table 1), for which genomic data and antibiotic susceptibility phenotype has previously been described^7,8^.

The second set, here-on-in described as ‘stainless steel biofilm analysis’ contained 260 *P. aeruginosa* genomes (Supp Table 2), 153 of which belong to the ST111 clone and 107 to the ST235 clone. Of these, 25 ST111 and 23 ST235 isolates were used to study biofilm phenotypes. All genomic data were downloaded from Genbank as fasta nucleotide sequences. Bacterial strains used for biofilm phenotyping were provided by bioMérieux (France) and Synthetic Genomics (USA) and were analysed in two previously published studies^6,7^.

### Construction of phylogenies

Core Single Nucleotide Polymorphisms (SNPs) were identified using the KSNP3 pipeline^24^. Kmer lengths of 21 were used for temperature-dependent biofilm analysis, whilst for stainless steel biofilm analysis, kmer lengths for ST111 and ST235 were 21 and 23 nucleotides respectively. Kmer lengths were selected using the KCHOOSER module within the KSNP3 suite. Bayesian phylogenetic analysis was performed using BEAST 2^25^. The following conditions were set: gamma site heterogeneity model, Hasegawa-Kishino-Yano (HKY) substitution model, relaxed-clock log-normal, chain length 5,000,000. BEAST 2 output was summarized using TreeAnnotator with a 5% burn in.

### Construction and interrogation of pan-genome

All genomes were annotated using PROKKA^26^. Using PROKKA output, pan-genomes were constructed using the Roary pan-genome pipeline^27^, and visualized with the interactive visualization tool Phandango ^28^.

### Biofilm phenotypes

Temperature-dependent biofilm analysis was carried out using standard non-modified 96-well plates (Starsted, Leicester, UK). Stainless steel biofilm analysis was performed on MBEC biofilm 96-well plates (Innovotech, Edmonton, Canada), with pegged lids spray coated with stainless steel. The sub-micron thick stainless steel coatings were deposited onto the lid by the physical vapour deposition technique of magnetron sputtering, which is widely used in industry to deposit thin (sub-micron to several micron thick) metallic or ceramic coatings onto a wide range of components^29^. The coatings were sputtered from a 300mm × 100mm 304 grade stainless steel target in a Teer Coatings Ltd UDP350 coating rig.

The method for biofilm growth was a modified version of that described by Coffey and Anderson^30^. *P. aeruginosa* strains were maintained on Luria-Bertani (LB) agar (BD, Sparks, MD) at 4°C. Liquid cultures were prepared by inoculating a single colony into 10 ml of LB broth (BD, Sparks, MD), and incubated at 37 °C with agitation (180 rpm). An overnight culture of each strain was diluted to an optical density (OD) of 1.0_540_. A 1:100 inoculation in sterile LB broth was made in a well of the MBEC plate base. Four wells per strain were inoculated. For temperature-dependent biofilm analysis, plates were incubated without shaking at either 22°C or 37°C for 48 hours.

For stainless steel biofilm analysis, the MBEC lid was replaced and the plate was incubated at 37°C with agitation (110 rpm) for 48 hours, with a change to a new 96-well plate with 200 μl of sterile LB broth in each well after 24 hours. To quantify biofilms, plates or pegs were gently immersed in water to remove non-adhered cells, placed in 0.1% crystal violet (Sigma, Dorset) for 10 minutes then gently washed in water, twice, to remove excess stain. The plates for temperature-dependent biofilm analysis had 200μl of 30% acetic acid added to each well to solubilize stain, whereas the lid with stained pegs was placed into of a 96-well plate containing 200μl per well of 30% acetic acid (Fisher Scientific, Loughborough) for 10 minutes with agitation (110 rpm). Optical density of each well was measured at 540 nm with a FLUOstar Omega plate reader (BMG Labtech, Aylesbury).

### Trait definition

In order to carry out GWAS and pan-GWAS (below), categorization of biofilm production was required. For temperature-dependent biofilm analysis, isolates were split into one of two traits: either producing more biofilm at 37°C compared to growth at 22°C, or producing more biofilm at 22°C compared to 37°C. For stainless steel biofilm analysis, a “good” biofilm formation trait was considered to be exhibited in strains providing an OD reading >= 0.5_540_, whilst poor biofilm formation was considered as an OD reading of < 0.5_540_.

### Pan-genome-wide association analysis

Scoary is a software tool able to provide pan-genome associations (pan-GWAS) based on user-defined traits using an input (gene presence / absence) file generated by Roary^31^. Analysis was performed independently for temperature-dependent biofilm traits as for both ST111 and ST235 genomes in the stainless steel biofilm analysis. Statistical significance of the associations was tested by Scoary with a Fisher’s exact test, and a pairwise comparisons algorithm to control spurious associations from stratified populations. *P* values were corrected for multiple comparisons using the Bonferroni adjustment. Where Scoary analysis provided genes significantly (*p*<0.05) associated with the biofilm phenotype that had not previously been annotated in PROKKA, and therefore considered hypothetical / putative, the amino acid sequence was searched in BLASTP, and if no hit was found, assessed using the sensitive protein homology detection, function and structure prediction tool HHPred (MPI Bioinformatics Tool Kit)^32^. The top three predictions were considered for assumed protein identity and function, assuming a match probability of at least 80%.

### SNP genome-wide association study

We used PLINK v1.90b5.3^33^ to explore SNP-based associations with user-defined traits. Analysis was performed in PLINK to test case / control associations with Fisher’s exact test, and return Bonferroni adjustments for statistical significance. Analysis was performed independently for temperature-dependent analysis as before for both ST111 and ST235 genomes in the stainless steel dataset using core SNP data produced by the KSNP3 pipeline. SNPs significantly associated with phenotype (*p*<0.05) where located in the genome using Artemis genome browser^34^. Genes containing SNPs of interest were interrogated using BLASTP and HHpred, with the top three predictions with a probability match of over 80% considered for assumed protein function.

## Results and discussion

### Temperature-dependent biofilm analysis

Biofilm formation varied greatly between isolates at both 22°C and 37°C (Figure 1). Of the 280 isolates studied, 69% (*n*=193) produced more biofilm at 22°C (average OD_540_ of 1.98) compared to 37°C (OD_540_ measurement of 1.29). When grown at 37°C, isolates demonstrated less variation in OD_540_ measures than at 22°C (Figure 2). Increased biofilm growth phenotype at 37°C was widely distributed across the phylogeny (Figure 3).

**Figure 1.**
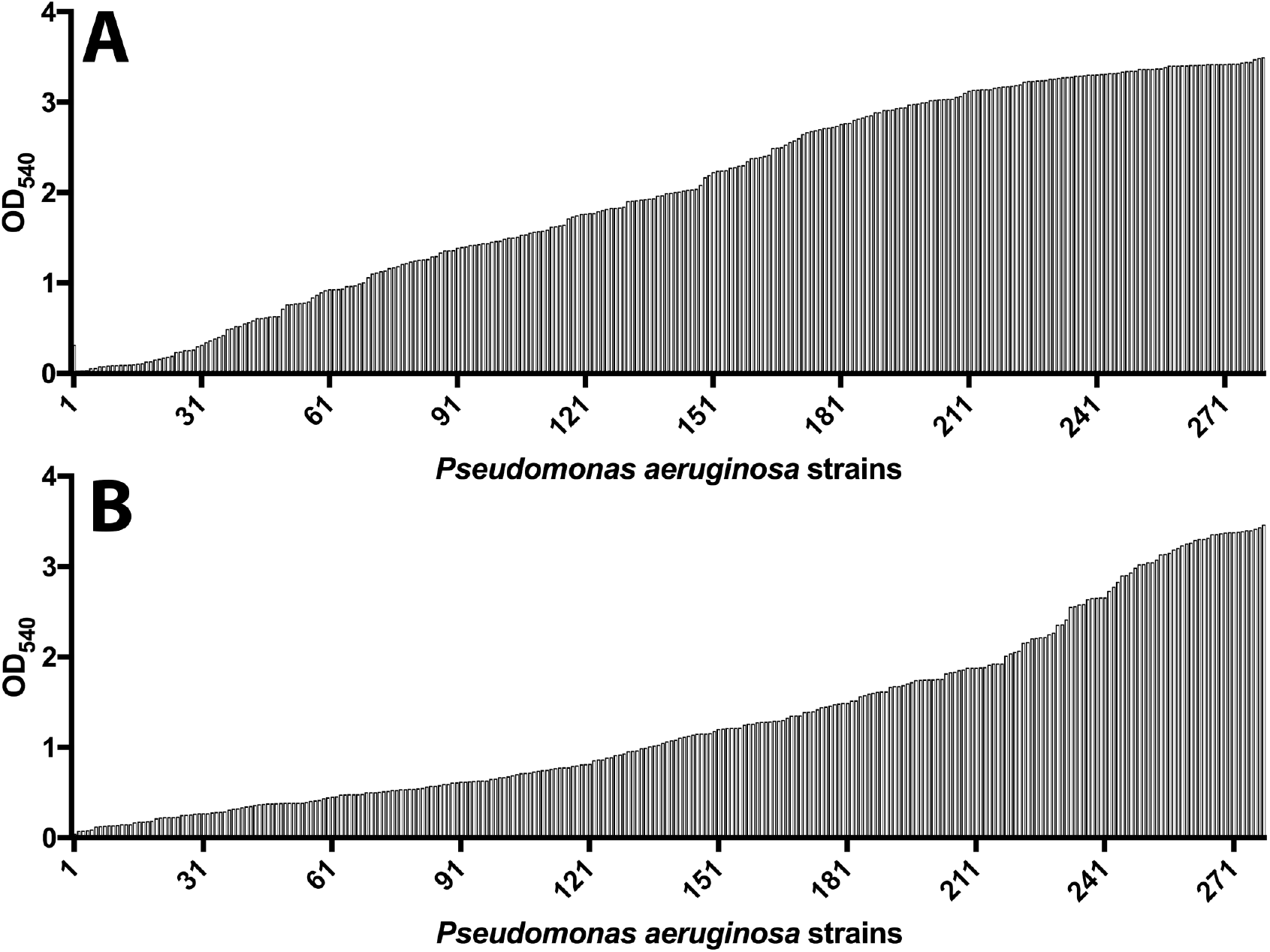
Optical density related to biofilm stained with 0.1% crystal violet and solubilised in 30% acetic acid at either 22 °C (A) or 37 °C (B) for 280 *Pseudomonas aeruginosa* isolates.

**Figure 2.**
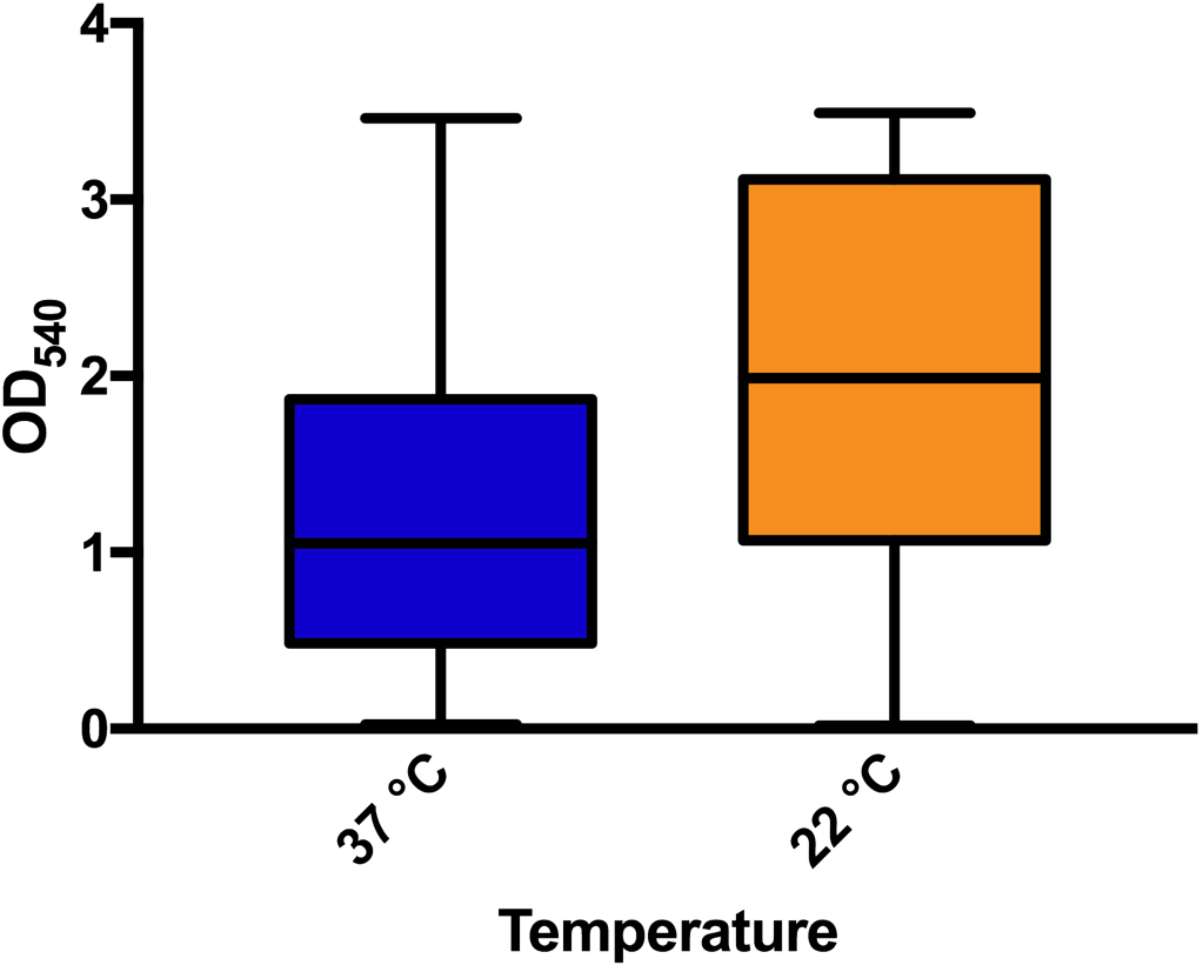
Box plot demonstrating the variation in biofilm formation for 280 *Pseudomonas aeruginosa* isolates at two different temperatures as measured by optical density.

**Figure 3.**
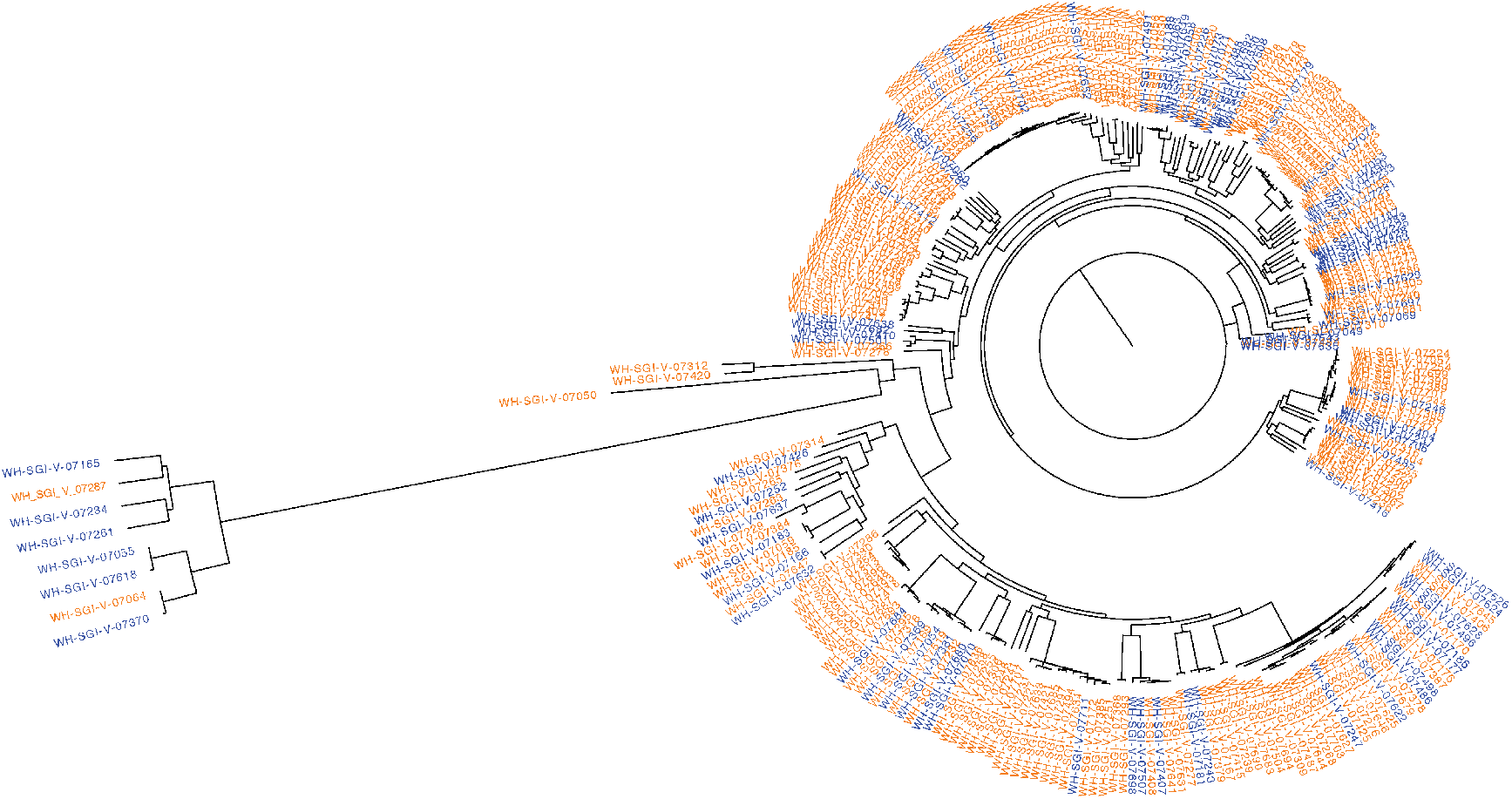
Maximum parsimony tree (from KSNP3 analysis) of 280 Pseudomonas aeruginosa isolates. Names highlighted orange represent an isolate that produced more biofilm at 22 °C and names highlighted blue represent an isolate that produced more biofilm at 37 °C.

Scoary analysis yielded 3,431 genes associated with the phenotypic trait of producing more biofilm at 22°C compared to 37°C, with only one gene passing with a significant Bonferoni *p* value (Table 1), annotated using BLASTN as *lpxO*, a lipid A hydroxylase gene. Lipid A is one of three distinct regions of the lipopolysaccharide of *P. aeruginosa*, and although literature on the gene is limited, it has been suggested that LpxO plays an essential part in the success of chronic infections (e.g. cystic fibrosis)^35,36^. More interestingly still, *lpxO* has been shown to be upregulated in *P. aeruginosa* when grown at a suboptimal temperature (22°C) compared to growth at 37°C^37^. Scoary analysis also provided other genes with significance, although with a Bonferonni *p* value of just above the 0.05 cut off (Table 1). These included *acr3, arsC, arsR* and *arsH*, four genes involved in arsenic resistance / reduction. Presence of arsenic in bacterial cells has been demonstrated to affect biofilm synthesis as well as chemotaxis and motility^38^, with suggestion that arsenic can prevent the switch between planktonic and sessile lifestyle^39^.

**Table 1.**
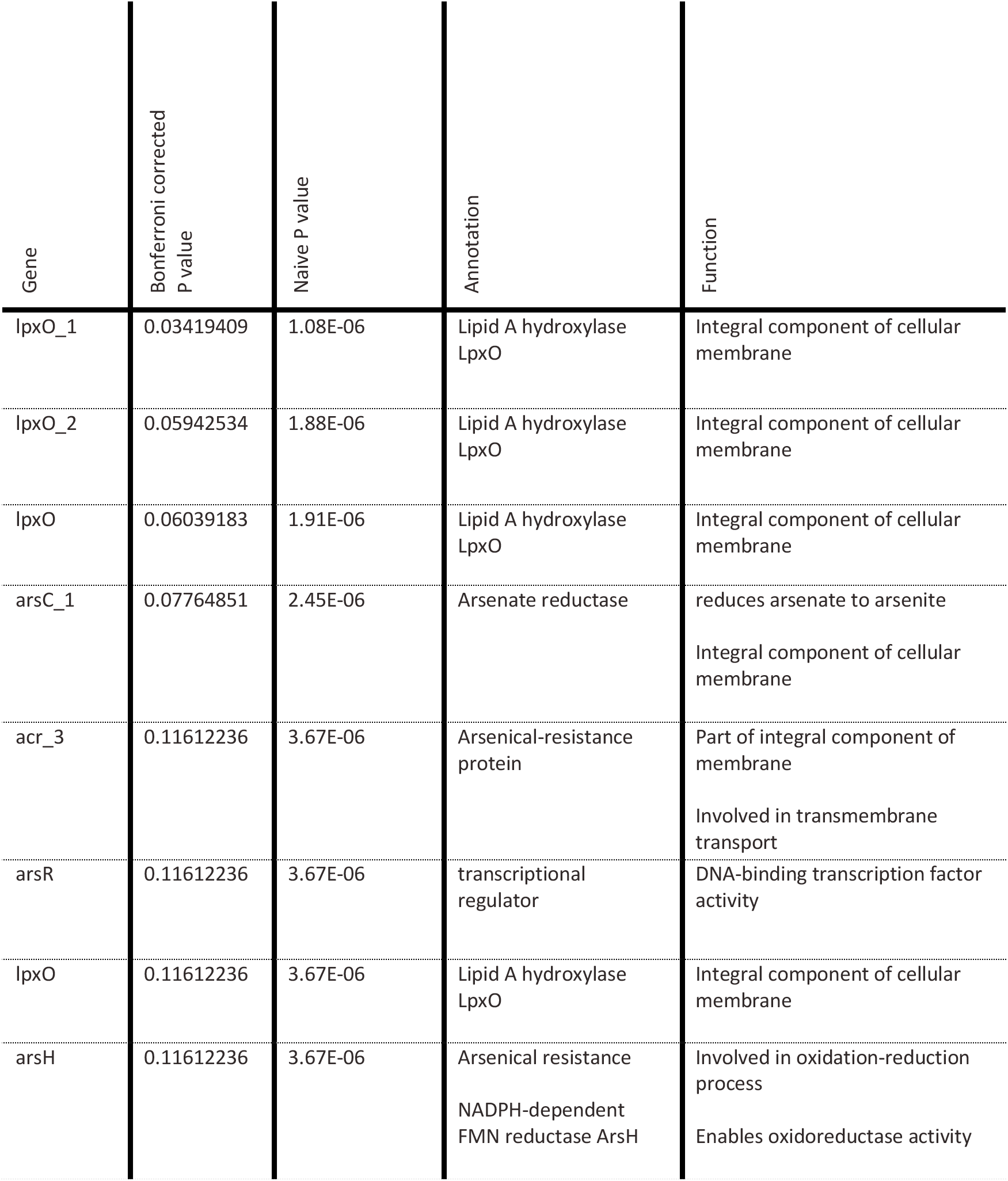
Genes associated with greater biofilm growth at 22 °C compared to 37 °C as determined by Scoary pan-genome-wide association analysis.

PLINK analysis yielded eleven significant SNPs (*p*<0.001 after Bonferroni adjustment) associated with increased biofilm production at 37°C compared to 22°C (Table 2). The most significant was a non-synonymous SNP found in the *lpxO* gene, the lipid A gene described earlier. Four other non-synonymous SNPs were identified as significant (*p*<0.001 after Bonferroni adjustment) and are present in a type 1 fimbrial gene, the *marC, pilW* and a hypothetical gene (*ydcV*). It is believed that type 1 fimbrial protein plays an essential role in the development of fimbrial structures, facilitiating bacterial attachement and promoting biofilm formation^40^. The *pilW* is involved in the development of the type-IV pilus, widely demonstrated as integral to twitching, swarming and biofilm production in *Pseudomonas* spp ^41,42^. *marC* encodes a multidrug resistance protein (homologous to *tspR*), deletion of which reduces type III secretion systems and motility whilst increasing biofilm production^43^. Of the remaining SNPs, four, whilst all present within a gene, were synonymous. One SNP was found in *arsH*, a gene which the Scoary gene presence/absence analysis also found to be of significant interest with the temperature-dependent biofilm phenotype (as described earlier). Another synonymous SNP of interest was located in a gene annotated as *smf-1*, a fimbriae-1 gene found in *Stenotrophomonas maltophilia*, which shares significant similarities to fimbrial adhesins found in the *cupA* fimbriae of *P. aeruginosa*, and has been demonstrated to be important in biofilm formation^44^.

**Table 2.**
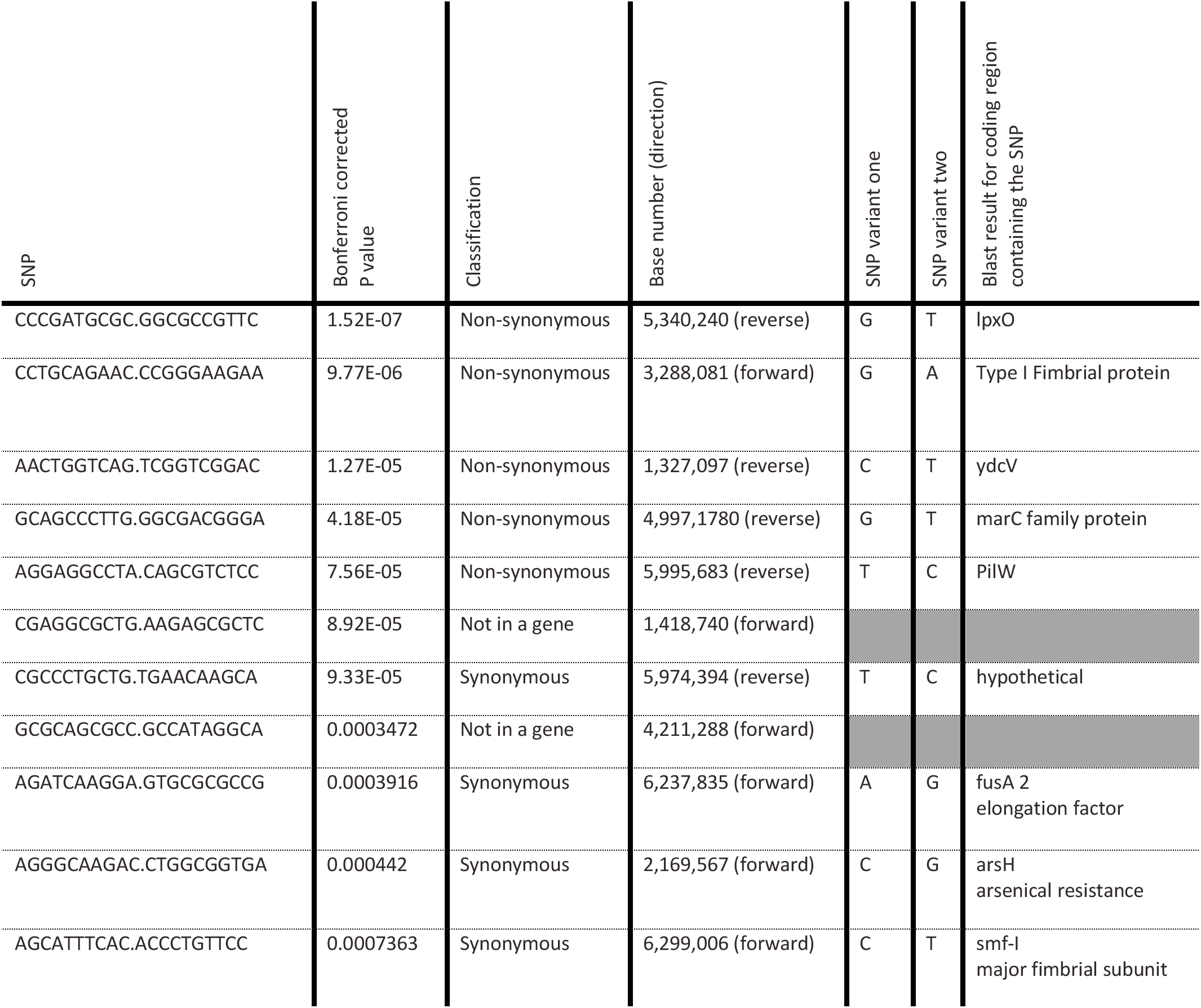
SNPs found statistically (p<0.05 using the Fisher exact test with Bonferroni adjustment) associated with more biofilm growth at 22 °C compared to 37 °C using PLINK. Base number relates to SNP position in isolate WH-SGI-V-07049.

### Biofilms on stainless steel

Of the isolates assessed for biofilm production on stainless steel (Figure 3), 40 % (*n*= 10 out of 25) ST111 and 52% (*n*=13 out of 23) ST235 were considered high biofilm producers. This demonstrates a large intra-clone variation of the biofilm phenotype. However, there was no significant difference between inter-clone variation, suggesting the intra-clone variation may be a common feature of other clones.

The BEAST 2 analysis suggests ST111 isolates diverged from a common ancestor ≅ 43 years ago (Figure 4.a), with a pan genome of 15,488 genes. The 107 ST235 isolates had a pan-genome of 15,178 genes and diverged from a common ancestor ≅ 28 years ago (Figure 4.b), suggesting that the two clones emerged within approximately 15 years of each other. When considered alongside biofilm phenotype, those isolates sharing phylogenetic similarity display similar biofilm phenotype, except for ST235 isolates WH-SGI-V-07622 and WH-SGI-V-07625, as well as isolates WH-SGI-V-07498 and WH-SGI-V-07624. Nevertheless, this analysis suggests many good and poor biofilm formers are closely related, and that biofilm production on stainless steel is not necessarily predictable based only on genomic analysis.

**Figure 4.**
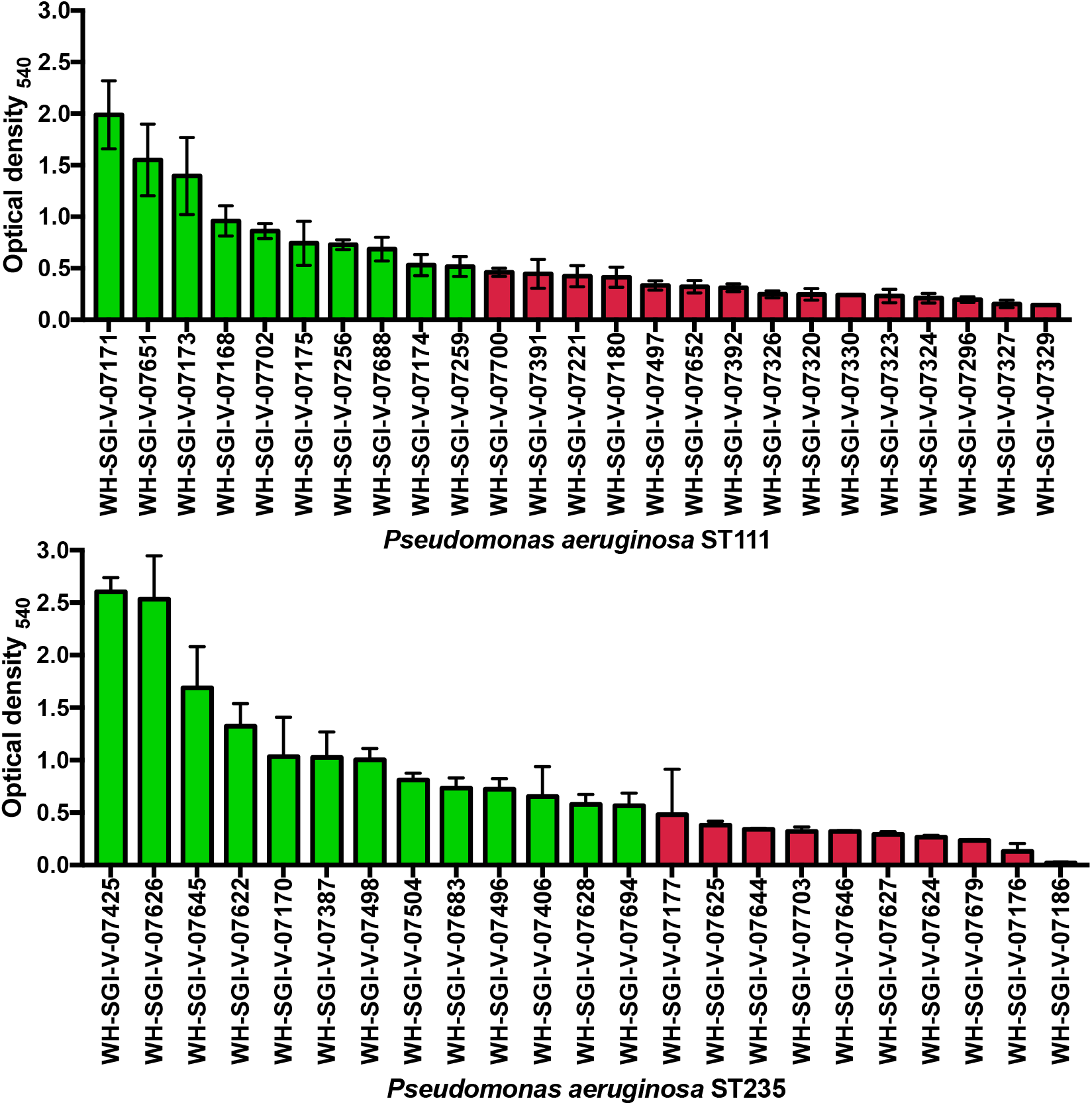
Optical density readings at 540nm of Pseudomonas aeruginosa biofilm after staining with 0.1% crystal violet and solubilised in 30% acetic acid. Biofilms were grown on a modified MBEC assay plate, of which the pegs had been coated in stainless steel. Data represent 25 ST111 (A) and 23 ST235 (B) strains. An optical density cut off of 0.5_540_ was used to differentiate high (green) and poor (red) biofilm producers.

The pan-genome of biofilm genes widely described in the literature (Figure 5) has a different core vs accessory structure in each of the clones, with the majority of genes (≈54%) in both clones present in less than 15% of genomes. ST235 has a higher percentage of core genes compared to ST111 (61.7% vs 31.34%), whilst ST111 has a larger cloud genome compared to ST235 (40.3% vs 17.02%). This variation is down to a number of homologous genes in the ST111 biofilm pan-genome *e.g. pelA, pslA, pslB*. The genes *pilA* and *fimT* were both identified in the ST111 pan-genome, but not found in the ST235 pan-genome, whilst *pslC* was found in ST235 but not in the ST111 pan-genome.

**Figure 5.**
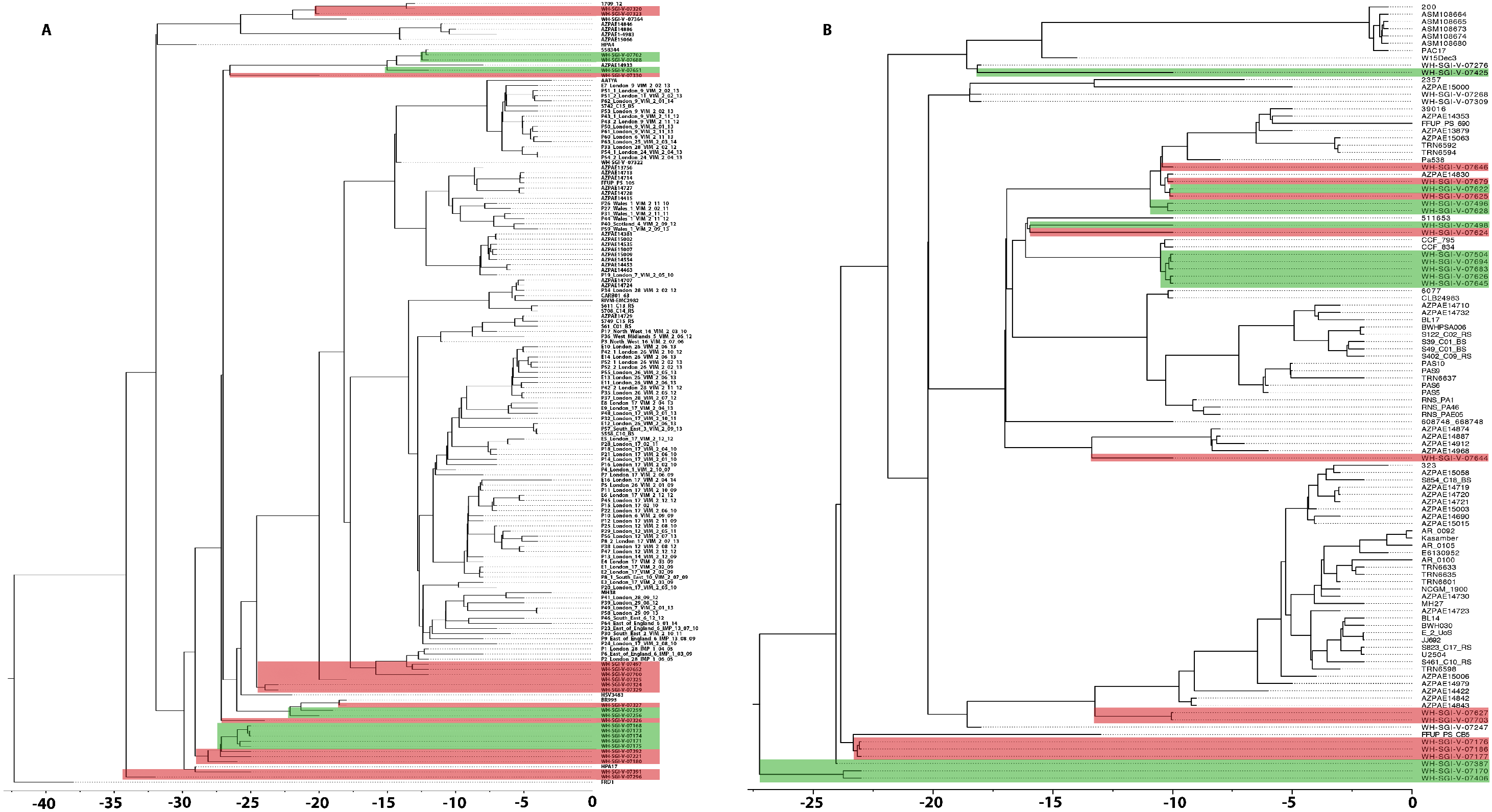
Bayesian evolutionary analysis sampling tree (BEAST 2) analysis representative of the ST111 (A) and ST235 (B) strains included within analysis. Horizontal distance is indicative of time in years, with 0 representing the most recent isolate (2017). Strains included in the biofilm on stainless steel phenotype assay (figure 4) are overlaid with their designated phenotype – high biofilm producers (green) and poor biofilm producers (red).

**Figure 6.**
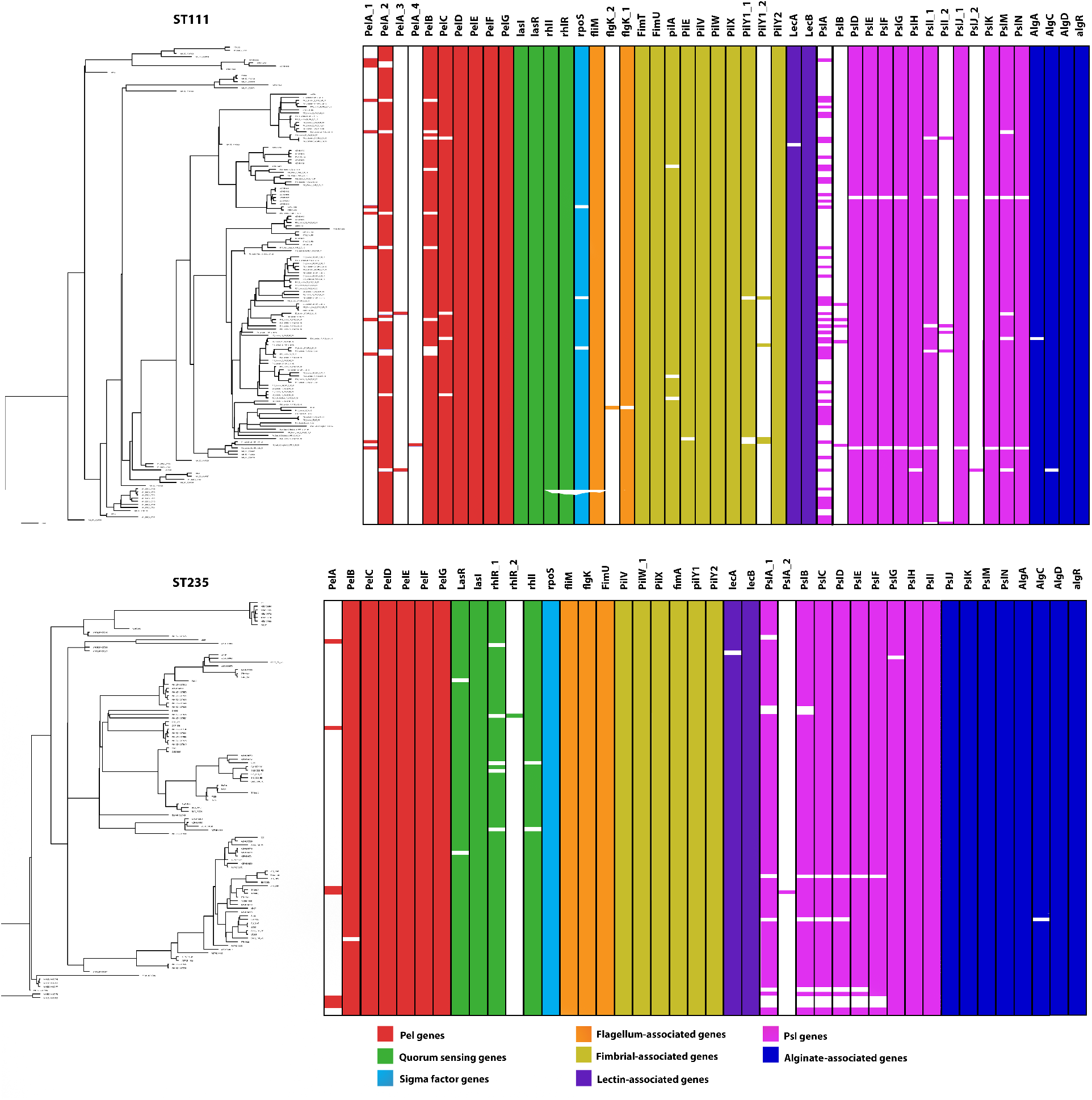
Biofilm gene presence/absence aligned with BEAST 2 analysis for ST111 (top) and ST235 (bottom) isolates (from Figure 2). Coloured blocks represent 48 ST111 and 42 ST235 genes that have previously been linked to biofilm formation in the literature, whilst white space represents gene absence. Colour of each block indicates gene families or biofilm features described in figure legend.

Scoary analysis provided 77 and 107 genes of interest for association with biofilm phenotype in ST111 and ST235 respectively although none had a Bonferonni of *p*<0.05. Of these genes, 24 ST111 and 30 ST235 genes did not provide a homology match of over 80% using HHpred, and would require further functional investigation to understand their potential association with biofilm phenotype. The remaining genes were considered for predicted function that might be obviously associated with biofilm formation (Table 3). ST111 had six genes with potential function associated with biofilm (7.8% of the ST111 Scoary output). Predicted homology of two of these genes suggest association of the flagellum system, while one gene shared 99.67% homology with *motB*, an essential gene of the *P. aeruginosa* flagellar motor. Mutants containing defects in the *mot* gene complex have been shown to exhibit a reduced or abolished motility^45^. Another gene shared 83.03% homology with *qseC*, a flagellum regulator gene with demonstrated importance in biofilm production in gram negative bacteria ^46,47^. Two further genes suggest association with the type-IV pili system,. These two genes share homology to a type-IV pili peptidoglycan endopeptidase (RipA – 92.55%) and a geopilin domain 1 protein (PilA – 92.32%). Of the two remaining ST111 genes from the Scoary analysis, one gene provided 100% homology to a subtilisin-like serine protease, a cellular mechanism which, if absent, has been shown to produce lower biofilm formation and motility function in *P. aeruginosa*^48^. The last of the ST111 Scoary genes shares 90.64% to BamC outer membrane protein, a lipoprotein likely involved in the outer-membrane biogenesis ^49^.

**Table 3.** Genes found statistically associated (unadjusted p<0.05) with biofilm phenotype using Scoary pan-genome-wide association analysis, for both ST111 or ST235.

| Gene ID | Predicted protein homology | Percentage similarity |
| --- | --- | --- |
| **ST111** |  |  |
| Group_6254 | Subtilisin-like serine protease | 100 |
| Group_8277 | PomB/MotB/ompA-like – involved in flagellar motor | 99.67 |
| Group_6216 | Type-IV pili peptidoglycan endopeptidase (RipA) | 92.55 |
| Group_1372 | Geopilin domain 1 protein; pilin, PilA, bacterial nanowire | 92.32 |
| Group_2185 | BamC outer membrane protein assembly factor | 90.64 |
| Group_8212 | Sensor protein qseC | 83.03 |
| **ST235** |  |  |
| Group_107 | Type-IV secretory hydrolase | 99.95 |
| Group_752 | Type-IV pili peptidoglycan endopeptidase (RipA) | 95.1 |
| Group_3943 | PilP protein | 98.12 |
| Group_859 | IcmL-like; type IV secretion system protein | 97.13 |
| Group_5868 | Flagellar transcriptional activator flhD/AcaD | 95.85 |

Of the ST235 Scoary analysis, five genes (4.7% of the ST235 Scoary output) had predicted function that might be obviously associated with biofilm formation (Table 3). Three of the genes share structural homology with type-IV secretion systems, two as hydrolases (one more specifically sharing homology with peptidoglycan endopeptidase RipA), and a third sharing homology to a type-IV secretion system-like gene IcmL-like,^50^. The fourth gene shared 98.12% homology with PilP, an essential part of the type-IV pili complex ^51^, whilst the fifth shared 95.85% homology with a flagellar transcriptional activator FlhD found in gram-negative bacteria ^52,53^. Whilst the Scoary pan-genome association analysis does not suggest an exact gene shared between the two clones in relation to the biofilm phenotype, it does emphasize the importance of the Type IV secretion systems (T4SS), with multiple genes present in both analyses. It is widely accepted and demonstrated in the literature that T4SS plays an important role in twitching motility (important for initial biofilm formation) as well as biofilm maturation^54,55^. It is acknowledged that biofilm production can be a highly specific trait, with environmental and mechanical changes and stresses altering the biofilm performance of a particular isolate, and so continued analysis of biofilm phenotypes in different models / environmental conditions would be beneficial.

PLINK analysis provided three SNPs of significant association with biofilm phenotype in ST235, and no significant SNPs in ST111 (Table 4). Cross checking SNPs with the relevant genome and BLASTP of the protein sequence shows that two of the SNPs were located in the same gene, a phage virion morphogeneisis protein, whilst the third was a non-coding SNP downstream of the viron morphogenesis protein. PaperBLAST^56^ searches provides on record of the gene or a homologue having already been explored in the literature.

**Table 4.** SNPs and related genes belonging to ST235 which have been found statistically (p<0.05 using the Fisher exact test with Bonferroni adjustment) associated with biofilm phenotype using PLINK genome-wide association analysis. Base number relates to SNP position in WH-SGI-V-07627.

| SNP | Bonferroni corrected<br>P value | Classification | Base number (direction) | SNP base for good biofilm<br>formers | SNP base for bad biofilm<br>formers | Amino acid coded for in good<br>biofilm formers | Amino acid coded for in good<br>biofilm formers | Blast result for coding region<br>containing the SNP |
| --- | --- | --- | --- | --- | --- | --- | --- | --- |
| CATGCTCA<br>GG.TGGAA<br>TGTGA | 0.006687 | Non-synonymous | 657,864 (f) | T | C | Threonine | Alanine | Pseudomonas<br>phage tail fiber<br>protein |
| GACGCCGC<br>TG.GCATA<br>ACGGA | 0.006687 | Synonymous | 657,937 (f) | C | T | Alanine | Alanine | Pseudomonas<br>phage tail fiber<br>protein |
| AGCGGCCA<br>CG.GTACAT<br>GCTC | 0.04681 | Noncoding | 657,850 (f) |  |  |  |  |  |

Whilst the phenotype and genotype analysis and associations described in this paper provide novel insight into the relationship between non-discreet phenotypes and genotype, standardization of such phenotypes are required. Currently, no agreed method exists for the categorization of good or poor biofilm formers, and much of the wider biofilm literature uses a dynamic range of methods and approaches to biofilm studies ^57^. This will be essential if GWAS and pan-GWAS studies are to be utilized to better understand the genetics of biofilm production.

## Conclusions

In this study we were able to identify candidate genes in which polymorphism (or deletion) was associated with differences in temperature-dependent biofilm phenotype for 280 *P. aeruginosa* isolates. The two different GWAS approaches used in this paper (SNP-based and gene presence / absence) both resulted in the identification of similar genes, namely *IpxO*, fimbrial genes and genes involved in arsenic resistance. The relationship between biofilm phenotypes on stainless steel and genotype in *P. aeruginosa* was less clear, likely due to the smaller subset of isolates used in this study. Further work should seek to utilize GWAS and pan-GWAS studies to identify new gene targets / cellular machinery alongside functional studies such gene expression (for example RNA-seq). Our study demonstrates the importance of understanding the complex nature of *P. aeruginosa* biofilm, and that phenotypic studies in isolation and/or assumptions that biofilm phenotype is consistent across *P. aeruginosa* offers little in the way of meaningful data. Unraveling the phenotype-genotype relationship of *P. aeruginosa* will be an essential next step in understanding and preventing biofilm in the era of AMR.

## References

1 Silby, M. W., Winstanley, C., Godfrey, S. A. C., Levy, S. B. & Jackson, R. W. Pseudomonas genomes: diverse and adaptable. FEMS Microbiology Reviews 35, 652–680, doi:10.1111/j.1574-6976.2011.00269.x (2011).

2 Boucher, H. W. et al. Bad bugs, no drugs: no ESKAPE ! An update from the Infectious Diseases Society of America. Clin Infect Dis 48, 1–12, doi:10.1086/595011 (2009).

3 Curran, B., Jonas, D., Grundmann, H., Pitt, T. & Dowson, C. G. Development of a multilocus sequence typing scheme for the opportunistic pathogen Pseudomonas aeruginosa. J Clin Microbiol 42, 5644–5649, doi:10.1128/JCM.42.12.5644-5649.2004 (2004).

4 Kos, V. N. et al. The resistome of Pseudomonas aeruginosa in relationship to phenotypic susceptibility. Antimicrob Agents Chemother 59, 427–436, doi:10.1128/AAC.03954-14 (2015).

5 Turton, J. F. et al. High-Resolution Analysis by Whole-Genome Sequencing of an International Lineage (Sequence Type 111) of Pseudomonas aeruginosa Associated with Metallo-Carbapenemases in the United Kingdom. Journal of Clinical Microbiology 53, 2622–2631, doi:10.1128/jcm.00505-15 (2015).

6 Jaillard, M. et al. Correlation between phenotypic antibiotic susceptibility and the resistome in Pseudomonas aeruginosa. Int J Antimicrob Agents 50, 210–218, doi:10.1016/j.ijantimicag.2017.02.026 (2017).

7 Alex van Belkum, L. B. S., Matthew C. LaFave, Srividya Akella, Jean-Baptiste Veyrieras, E. Magda Barbu, Dee Shortridge, Bernadette Blanc, Gregory Hannum, Gilles Zambardi, Kristofer Miller, Mark C. Enright, Nathalie Mugnier, Daniel Brami, Stéphane Schicklin, Martina Felderman, Ariel S. Schwartz, Toby H. Richardson, Todd C. Peterson, Bolyn Hubby, Kyle C. Cady. Phylogenetic Distribution of CRISPR-Cas Systems in Antibiotic-Resistant Pseudomonas aeruginosa. mbio 6, e01796–01715 (2015).

8 Pirnay, J.-P. et al. Pseudomonas aeruginosa Population Structure Revisited. PLOS ONE 4, e7740, doi:10.1371/journal.pone.0007740 (2009).

9 Oliver, A., Mulet, X., López-Causapé, C. & Juan, C. The increasing threat of Pseudomonas aeruginosa high-risk clones. Drug Resistance Updates 21-22, 41–59, doi:https://doi.org/10.1016/j.drup.2015.08.002 (2015).

10 Sánchez-Diener, I. et al. Interplay among Resistance Profiles, High-Risk Clones, and Virulence in the Caenorhabditis elegans Pseudomonas aeruginosa Infection Model. Antimicrobial Agents and Chemotherapy 61, doi:10.1128/aac.01586-17 (2017).

11 Treepong, P. et al. Global emergence of the widespread Pseudomonas aeruginosa ST235 clone. Clinical Microbiology and Infection 24, 258–266, doi:10.1016/j.cmi.2017.06.018 (2018).

12 Quainoo, S. et al. Whole-Genome Sequencing of Bacterial Pathogens: the Future of Nosocomial Outbreak Analysis. Clinical Microbiology Reviews 30, 1015–1063, doi:10.1128/cmr.00016-17 (2017).

13 Aanensen, D. M. et al. Whole-Genome Sequencing for Routine Pathogen Surveillance in Public Health: a Population Snapshot of Invasive Staphylococcus aureus in Europe. mBio 7, doi:10.1128/mBio.00444-16 (2016).

14 Rasamiravaka, T., Labtani, Q., Duez, P. & El Jaziri, M. The Formation of Biofilms by Pseudomonas aeruginosa: A Review of the Natural and Synthetic Compounds Interfering with Control Mechanisms. BioMed Research International 2015, 17, doi:10.1155/2015/759348 (2015).

15 Hutchison, M. L. & Govan, J. R. W. Pathogenicity of microbes associated with cystic fibrosis. Microbes and Infection 1, 1005–1014, doi:https://doi.org/10.1016/S1286-4579(99)80518-8 (1999).

16 Wendel, A. F., Ressina, S., Kolbe-Busch, S., Pfeffer, K. & MacKenzie, C. R. Species Diversity of Environmental GIM-1-Producing Bacteria Collected during a Long-Term Outbreak. Applied and Environmental Microbiology 82, 3605–3610, doi:10.1128/aem.00424-16 (2016).

17 Verran, J. & Redfern, J. in Handbook of Hygiene Control in the Food Industry Vol. 2 (eds Huub Lelieveld, John Holah, & Domagoj Gabrić) Ch. 42, 651–659 (Woodhead Publishing Limited, 2016).

18 Salm, F. et al. Prolonged outbreak of clonal MDR Pseudomonas aeruginosa on an intensive care unit: contaminated sinks and contamination of ultra-filtrate bags as possible route of transmission? Antimicrobial Resistance & Infection Control 5, 53, doi:10.1186/s13756-016-0157-9 (2016).

19 Mulet, X. et al. Biological Markers of Pseudomonas aeruginosa Epidemic High-Risk Clones. Antimicrobial Agents and Chemotherapy 57, 5527–5535, doi:10.1128/aac.01481-13 (2013).

20 Varin, A. et al. High prevalence and moderate diversity of Pseudomonas aeruginosa in the U-bends of high-risk units in hospital. International Journal of Hygiene and Environmental Health 220, 880–885, doi:http://dx.doi.org/10.1016/j.ijheh.2017.04.003 (2017).

21 Gbaguidi-Haore, H. et al. A Bundle of Measures to Control an Outbreak of Pseudomonas aeruginosa Associated With P-Trap Contamination. Infection Control & Hospital Epidemiology 39, 164–169, doi:10.1017/ice.2017.304 (2018).

22 Chen, P. E. & Shapiro, B. J. The advent of genome-wide association studies for bacteria. Current Opinion in Microbiology 25, 17–24, doi:https://doi.org/10.1016/j.mib.2015.03.002 (2015).

23 Read, T. D. & Massey, R. C. Characterizing the genetic basis of bacterial phenotypes using genome-wide association studies: a new direction for bacteriology. Genome medicine 6, 109–109, doi:10.1186/s13073-014-0109-z (2014).

24 Gardner, S. N., Slezak, T. & Hall, B. G. kSNP3.0: SNP detection and phylogenetic analysis of genomes without genome alignment or reference genome. Bioinformatics 31, 2877–2878, doi:10.1093/bioinformatics/btv271 (2015).

25 Bouckaert, R. et al. BEAST 2: A Software Platform for Bayesian Evolutionary Analysis. PLOS Computational Biology 10, e1003537, doi:10.1371/journal.pcbi.1003537 (2014).

26 Seemann, T. Prokka: rapid prokaryotic genome annotation. Bioinformatics 30, 2068–2069, doi:10.1093/bioinformatics/btu153 (2014).

27 Page, A. J. et al. Roary: rapid large-scale prokaryote pan genome analysis. Bioinformatics 31, 3691–3693, doi:10.1093/bioinformatics/btv421 (2015).

28 Hadfield, J. et al. Phandango: an interactive viewer for bacterial population genomics. Bioinformatics, btx610–btx610, doi:10.1093/bioinformatics/btx610 (2017).

29 Kelly, P. J. & Arnell, R. D. Magnetron sputtering: a review of recent developments and applications. Vacuum 56, 159–172, doi:https://doi.org/10.1016/S0042-207X(99)00189-X (2000).

30 Coffey, B. M. & Anderson, G. G. in Pseudomonas Methods and Protocols (eds Alain Filloux & Juan-Luis Ramos) 631–641 (Springer New York, 2014).

31 Brynildsrud, O., Bohlin, J., Scheffer, L. & Eldholm, V. Rapid scoring of genes in microbial pan-genome-wide association studies with Scoary. Genome Biology 17, 238, doi:10.1186/s13059-016-1108-8 (2016).

32 Zimmermann, L. et al. A Completely Reimplemented MPI Bioinformatics Toolkit with a New HHpred Server at its Core. Journal of Molecular Biology, doi:https://doi.org/10.1016/j.jmb.2017.12.007 (2017).

33 Purcell, S. et al. PLINK: A Tool Set for Whole-Genome Association and Population-Based Linkage Analyses. American Journal of Human Genetics 81, 559–575 (2007).

34 Carver, T., Harris, S. R., Berriman, M., Parkhill, J. & McQuillan, J. A. Artemis: an integrated platform for visualization and analysis of high-throughput sequence-based experimental data. Bioinformatics 28, 464–469, doi:10.1093/bioinformatics/btr703 (2012).

35 Moskowitz, S. M. & Ernst, R. K. The role of Pseudomonas lipopolysaccharide in cystic fibrosis airway infection. Sub-cellular biochemistry 53, 241–253, doi:10.1007/978-90-481-9078-2_11 (2010).

36 Lam, J., Taylor, V., Islam, S., Hao, Y. & Kocíncová, D. Genetic and Functional Diversity of Pseudomonas aeruginosa Lipopolysaccharide. Frontiers in microbiology 2, doi:10.3389/fmicb.2011.00118 (2011).

37 Termi, F. & Michel, G. P. Transcriptome and secretome analyses of the adaptive response of Pseudomonas aeruginosa to suboptimal growth temperature. International Microbiology 12, 7 (2009).

38 Andres, J. & Bertin, P. N. The microbial genomics of arsenic. FEMS Microbiology Reviews 40, 299–322, doi:10.1093/femsre/fuv050 (2016).

39 Marchal, M., Briandet, R., Koechler, S., Kammerer, B. & Bertin, P. N. Effect of arsenite on swimming motility delays surface colonization in Herminiimonas arsenicoxydans. Microbiology 156, 2336–2342, doi:doi:10.1099/mic.0.039313-0 (2010).

40 Ruer, S., Stender, S., Filloux, A. & de Bentzmann, S. Assembly of Fimbrial Structures in <em>Pseudomonas aeruginosa</em>: Functionality and Specificity of Chaperone-Usher Machineries. Journal of Bacteriology 189, 3547–3555, doi:10.1128/jb.00093-07 (2007).

41 Jarrell, K. F. & Albers, S.-V. The archaellum: an old motility structure with a new name. Trends in Microbiology 20, 307–312, doi:https://doi.org/10.1016/j.tim.2012.04.007 (2012).

42 Otton, L. M., da Silva Campos, M., Meneghetti, K. L. & Corção, G. Influence of twitching and swarming motilities on biofilm formation in Pseudomonas strains. Archives of Microbiology 199, 677–682, doi:10.1007/s00203-017-1344-7 (2017).

43 Zhu, M. et al. Modulation of Type III Secretion System in Pseudomonas aeruginosa: Involvement of the PA4857 Gene Product. Frontiers in microbiology 7, doi:10.3389/fmicb.2016.00007 (2016).

44 De Oliveira-Garcia, D. et al. Fimbriae and adherence of Stenotrophomonas maltophilia to epithelial cells and to abiotic surfaces. Cellular Microbiology 5, 625–636, doi:10.1046/j.1462-5822.2003.00306.x (2003).

45 Doyle, T. B., Hawkins, A. C. & McCarter, L. L. The Complex Flagellar Torque Generator of Pseudomonas aeruginosa. Journal of Bacteriology 186, 6341–6350, doi:10.1128/jb.186.19.6341-6350.2004 (2004).

46 Curtis, M. M. et al. QseC Inhibitors as an Antivirulence Approach for Gram-Negative Pathogens. mBio 5, doi:10.1128/mBio.02165-14 (2014).

47 Novak, E. A., Shao, H., Daep, C. A. & Demuth, D. R. Autoinducer-2 and QseC Control Biofilm Formation and In Vivo Virulence of Aggregatibacter actinomycetemcomitans. Infection and immunity 78, 2919–2926, doi:10.1128/iai.01376-09 (2010).

48 Fernández, L., Breidenstein, E. B. M., Song, D. & Hancock, R. E. W. Role of Intracellular Proteases in the Antibiotic Resistance, Motility, and Biofilm Formation of Pseudomonas aeruginosa. Antimicrobial Agents and Chemotherapy 56, 1128–1132, doi:10.1128/AAC.05336-11 (2012).

49 Remans, K., Vercammen, K., Bodilis, J. & Cornelis, P. Genome-wide analysis and literature-based survey of lipoproteins in Pseudomonas aeruginosa. Microbiology 156, 2597–2607, doi:doi:10.1099/mic.0.040659-0 (2010).

50 Segal, G., Feldman, M. & Zusman, T. The Icm/Dot type-IV secretion systems of Legionella pneumophila and Coxiella burnetii. FEMS Microbiology Reviews 29, 65–81, doi:10.1016/j.femsre.2004.07.001 (2005).

51 Burrows, L. L. Pseudomonas aeruginosa Twitching Motility: Type IV Pili in Action. Annual Review of Microbiology 66, 493–520, doi:10.1146/annurev-micro-092611-150055 (2012).

52 Prüß, B. M., Liu, X., Hendrickson, W. & Matsumura, P. FlhD/FlhC – regulated promoters analyzed by gene array and lacZ gene fusions. FEMS Microbiology Letters 197, 91–97, doi:doi:10.1111/j.1574-6968.2001.tb10588.x (2001).

53 Liu, X. & Matsumura, P. The FlhD/FlhC complex, a transcriptional activator of the Escherichia coli flagellar class II operons. Journal of Bacteriology 176, 7345–7351, doi:10.1128/jb.176.23.7345-7351.1994 (1994).

54 O’Toole, G. A. & Kolter, R. Flagellar and twitching motility are necessary for Pseudomonas aeruginosa biofilm development. Molecular Microbiology 30, 295–304, doi:doi:10.1046/j.1365-2958.1998.01062.x (1998).

55 Barken, K. B. et al. Roles of type IV pili, flagellum-mediated motility and extracellular DNA in the formation of mature multicellular structures in Pseudomonas aeruginosa biofilms. Environmental Microbiology 10, 2331–2343, doi:doi:10.1111/j.1462-2920.2008.01658.x (2008).

56 Price, M. N. & Arkin, A. P. PaperBLAST: Text Mining Papers for Information about Homologs. mSystems 2, doi:10.1128/mSystems.00039-17 (2017).

57 Coenye, T., Goeres, D., Van Bambeke, F. & Bjarnsholt, T. Should standardized susceptibility testing for microbial biofilms be introduced in clinical practice? Clinical Microbiology and Infection 24, 570–572, doi:https://doi.org/10.1016/j.cmi.2018.01.003 (2018).

